# Detection of Viral RNA Fragments in Human iPSC-Cardiomyocytes following Treatment with Extracellular Vesicles from SARS-CoV-2 Coding-Sequence-Overexpressing Lung Epithelial Cells

**DOI:** 10.1101/2020.05.14.093583

**Authors:** Youjeong Kwon, Sarath Babu Nukala, Shubhi Srivastava, Hiroe Miyamoto, Nur Izzah Ismail, Jalees Rehman, Sang-Bing Ong, Won Hee Lee, Sang-Ging Ong

## Abstract

The novel coronavirus disease 2019 (COVID-19) caused by the severe acute respiratory syndrome coronavirus 2 (SARS-CoV-2) has evolved into a worldwide pandemic. Early data suggest that the prevalence and severity of COVID-19 appear to be higher among patients with underlying cardiovascular risk factors. Despite the expression of angiotensin-converting enzyme 2 (ACE2), a functional receptor for SARS-CoV-2 infection, in cardiomyocytes, there has been no conclusive evidence of direct viral infection although the presence of inflammation and viral genome within the hearts of COVID-19 patients have been reported. Here we transduced A549 lung epithelial cells with lentivirus overexpressing selected genes of the SARS-CoV-2. We then isolated extracellular vesicles (EVs) from the supernatant of A549 cells and detected the presence of viral RNA within the purified EVs. Importantly, we observed that human induced pluripotent stem cell-derived cardiomyocytes (hiPSC-CMs) were able to actively uptake these EVs and viral genes were subsequently detected in the cardiomyocytes. Accordingly, uptake of EVs containing viral genes led to an upregulation of inflammation-related genes in hiPSC-CMs. Thus, our findings indicate that SARS-CoV-2 RNA-containing EVs represent an indirect route of viral RNA entry into cardiomyocytes.

## INTRODUCTION

Since the initial outbreak in China, coronavirus disease 2019 (COVID-19) which is caused by severe acute respiratory syndrome coronavirus 2 (SARS-CoV-2) has evolved into a global pandemic. While COVID-19 affects both healthy individuals as well as those with comorbid conditions such as cardiovascular diseases, the severity and risk of adverse outcomes of COVID-19 are especially pronounced in the latter^1^. Furthermore, patients with COVID-19 have also been reported to exhibit increased levels of cardiac biomarkers, suggestive of cardiac injury^2^. However, it remains unclear whether exacerbated cardiac injury seen in COVID-19 patients results directly from viral SARS-CoV-2 infection of the myocardium or indirectly from the complications of COVID-19. Cardiomyocytes express angiotensin-converting enzyme 2 (ACE2), the SARS-CoV-2 binding site^3^. However, there is no evidence of direct viral infection of cardiomyocytes to date, although the presence of myocardial inflammation and viral particles among the interstitial cells of the myocardium have been reported^4^ and viral RNA has also detected in some COVID-19 patients’ hearts^5^. The majority of cells in the body are known to release lipid bilayer membrane vesicles, also known as extracellular vesicles (EVs), that are capable of transferring various genetic materials including viral RNAs to other recipient cells^6^, ^7^. Therefore, in the present work, we hypothesized that SARS-CoV-2-infected cells such as airway epithelial cells secrete EVs carrying viral genetic material that may be taken up by cardiomyocytes and establish an indirect route of SARS-CoV-2 genetic material transmission.

## RESULTS AND DISCUSSION

To test whether the viral RNA of SARS-CoV-2 can be transmitted via EVs into cardiomyocytes without the need for direct infection, we transduced A549 lung epithelial cells with lentivirus encoding selected SARS-CoV-2 proteins^8^ (**Figure 1A**). A549 cells were chosen as a model cell type since COVID-19 appears to mainly infect respiratory tract cells in patients. SARS-CoV-2 genes encoding for two non-structural proteins (*Nsp1* and *Nsp12*) and two structural proteins (envelope *E* and nucleocapsid *N*) were used for this proof-of-principle study. We opted not to include the spike (*S*) protein, which is required for receptor binding and viral entry, in order to focus on EV-mediated transfer of viral fragments into recipient cardiomyocytes that is independent of S-protein mediated direct viral entry. The use of lentivirus overexpressing viral subunits also allowed us to distinguish EV-mediated SARS-CoV-2 RNA transfer from canonical virus infection since EV preparations inevitably contain infectious virions due to the overlap in size.

**Figure 1.**
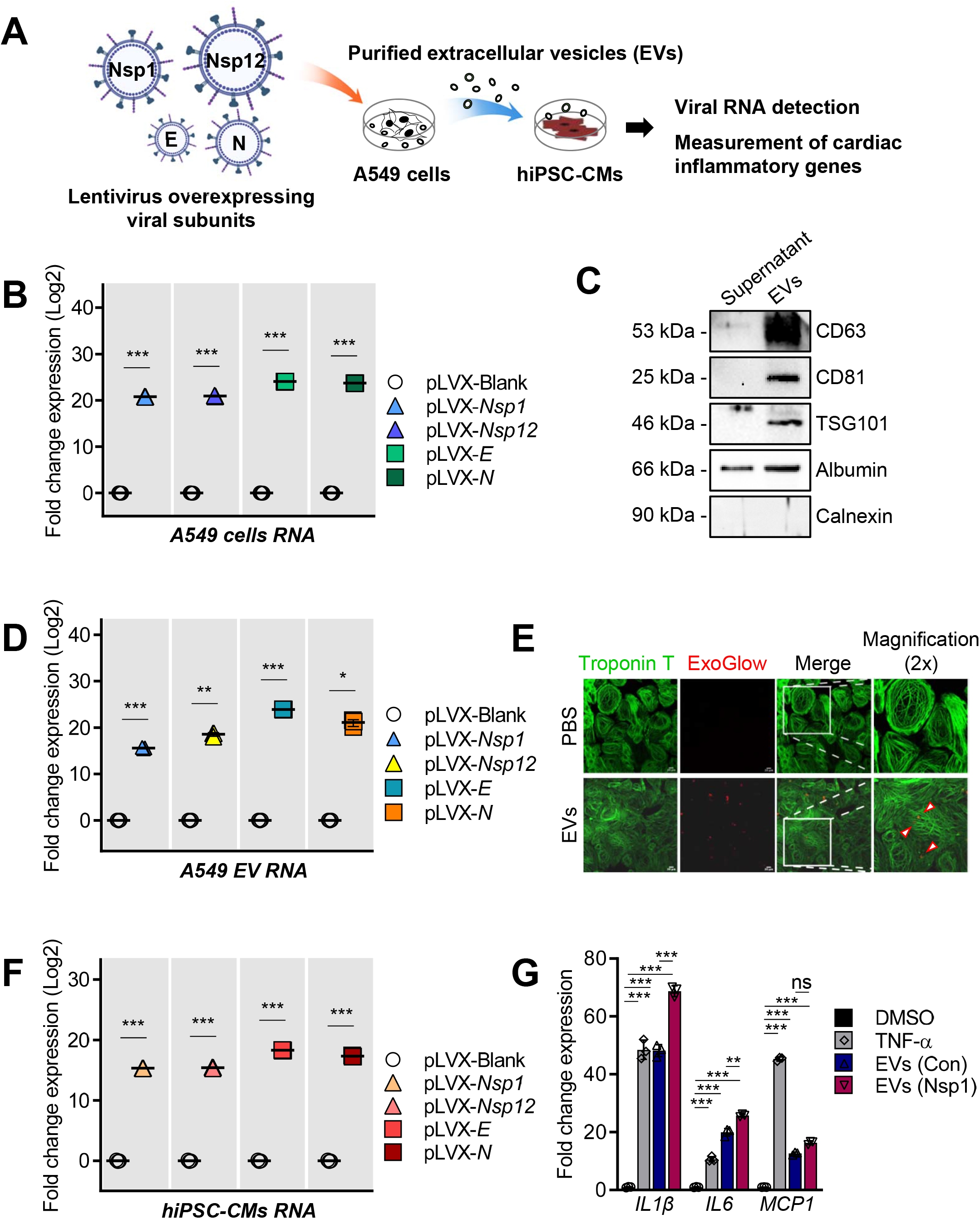
Transmission of SARS-CoV-2 synthetic viral RNA into human induced pluripotent stem cell-derived cardiomyocytes (hiPSC-CMs) through EVs. **A**, Schematic depiction of study design. Nsp1 indicates non-structural protein 1; Nsp12, non-structural protein 12; E, envelope protein; N, nucleocapsid protein. **B**, Expression of SARS-Cov-2 genes in A549 lung epithelial cells. A549 cells were infected with indicated lentiviral particles for 48 hours and mRNA levels were measured by qRT-PCR (n=3, mean± S.D). ***P<0.001 versus pLVX-Blank (Student’s t-test). **C**, Immunoblotting of EV markers demonstrating enrichment in the EV fraction compared to supernatant. **D**, SARS-CoV-2 genetic materials (Nsp1, Nsp12, E and N) were detected in EVs secreted from A549 lung epithelial cells. EVs were purified from A549 cell culture media and mRNA levels were measured by qRT-PCR (n = 3, mean ± S.D.). *P<0.05; **P<0.01; ***P<0.001 versus pLVX-Blank (Student’s t-test). **E**, Uptake of ExoGlow-labeled EVs (pseudocolored red) or PBS (negative control) by hiPSC-CMs stained with cardiac troponin T (green) were visualized by confocal imaging. Scale bar = 10 μM. Small arrows depict detected EVs within cells. **F**, qRT-PCR was performed to detect the presence of viral genes in hiPSC-CMs following EV uptake. mRNA levels were measured by qRT-PCR (n = 3, mean ± S.D.). ***P<0.001 versus pLVX-Blank (Student’s t-test). **G**, Expression of inflammatory genes in hiPSC-CMs. hiPSC-CMs were treated with EVs released by A549 cells transduced with pLVX-Blank or pLVX-Nsp1 lentiviral particles for 6 hours and mRNA levels were measured by qRT-PCR (n = 3, mean ± S.D.). Tumor necrosis factor-α (TNF-α, 50 ng/ml) was used as a positive control. **P<0.01; ***P<0.001; n.s., not significant (Two-way ANOVA followed by Tukey’s multiple comparisons test). Interleukin 1β (*IL1β);* interleukin 6 (*IL6*); monocyte chemoattractant protein 1 (*MCP1*).

Quantitative RT-PCR on total RNA extracted from A549 cells 48 hours after lentivirus transduction confirmed the successful overexpression of viral RNAs encoding for Nsp1, Nsp12, E and N respectively compared to a control empty vector (**Figure 1B**). To isolate EVs released by A549 cells, the supernatant of A549 cells grown in culture medium supplemented with exosome-depleted FBS for 48 hours was collected for EV purification. Immunoblotting of EV preparations confirmed the enrichment of the EV markers CD63, CD81 and TSG101 although we did note the presence of albumin most likely due to the PEG-based isolation method used (**Figure 1C**). In addition, there was no significant difference between control and viral proteins overexpressing A549 cells in terms of EVs production with an average of 100 μg of EVs being secreted over a period of 48 hours by 1 x 10^7^ cells (data not shown).

We next asked whether the RNAs encoding for SARS-CoV-2 are packaged into purified EVs of A549 cells. qRT-PCR revealed the presence of mRNA in purified EVs for each of the four tested SARS-CoV-2 genes (**Figure 1D**). We performed a separate validation for Nsp1 and Nsp12 in EVs isolated using a different method based on immuno-magnetic CD63-beads and successfully confirmed the presence of both tested genes in the isolated EVs (**Figure S1**). To further substantiate the presence of viral genes within EVs, we treated our Nsp1 EV preparation with RNase/protease. As expected, treatment with RNase alone or protease + RNase led to minimal loss of Nsp1, while addition of detergent led to significant degradation of the enclosed Nsp1 (**Figure S2**). To study if human cardiomyocytes are able to uptake EVs, we labeled EVs with a fluorescent dye ExoGlow, and incubated them (100 μg) with human induced pluripotent stem cells-derived cardiomyocytes (hiPSC-CMs, 1 x 10^6^ cells). Following 6 hours of 37°C incubation and washout of unbound EVs, we observed the presence of labeled EVs in treated hiPSC-CMs, which was not observed when the cells were incubated with the negative control (PBS without EVs stained with ExoGlow), confirming the successful binding/uptake of EVs by the recipient hiPSC-CMs (**Figure 1E** and **Figure S3**). Moreover, after exposure of hiPSC-CMs to A549 EVs for 24 hours, we detected all four tested viral RNAs in the hiPSC-CMs, but we could not detect any significant levels of viral RNAs in hiPSC-CMs treated with control EVs (**Figure 1F**). In a separate set of experiments, we exposed hiPSC-CMs to conditioned media collected from Nsp1-overexpressing A549 cells (1 x 10^7^ cells) with and without concurrent treatment of GW4869 (5 μM), an inhibitor of exosome generation. The expression of Nsp1 in hiPSC-CMs was significantly blunted when GW4869 was present, consistent with the involvement of EVs in RNA transfer (**Figure S4**). We next assessed whether hiPSC-CM exposure to EVs from A549 cells expressing viral RNA increased inflammatory gene expression. It is known that, A549 EVs themselves can increase inflammation^9^. We noted that EVs containing Nsp1 further increased the expression of the pro-inflammatory genes *IL1β* and *IL6,* suggesting the transferred viral gene may promote inflammation (**Figure 1G**).

There are several limitations in our study. Although the precipitation method used in our study for isolating EVs leads to high recovery, it is associated with recovery of non-EV components such as proteins. However, our additional experiments including immuno-magnetic isolation of EVs, RNase/protease treatment and GW4869 treatment support the enrichment of viral genes within EVs. The ExoGlow dye used cannot fully distinguish among binding and internalization of EVs. It should also be noted that the viral genes used in our study are codon-optimized and expressed at a supraphysiological level. Further experiments are needed to validate if EVs are capable of transferring actual viral RNAs or virions at a physiological level, and if the transferred RNAs are biologically active. Overall, our results collectively demonstrated that lung epithelial cells expressing SARS-CoV-2 genes can secrete EVs containing viral RNA fragments that can be detected in cardiomyocytes suggesting an indirect route of viral RNA delivery into cardiac cells via EVs. Transfer of viral RNA via EVs should be considered when studying the widespread multi-organ effects of a SARS-CoV-2 infection that has been reported^10^, because it indicates that cells which do not express the SARS-CoV-2 receptor ACE2 might still be vulnerable via the uptake of EVs. Further work is needed to clarify whether the entry of SARS-CoV-2 RNA via EVs is sufficient to induce cell injury and inflammation.

## METHODS

### Cell Culture

HEK293T cells were purchased from TakaraBio and cultured in Dulbecco’s modified Eagle’s medium (DMEM) with high glucose (Thermo Fisher, 11995-065) supplemented with 10% serum. A549 cells were purchased from ATCC and cultured in Ham’s F-12K medium (Thermo Fisher, 21127022). Human induced pluripotent stem cells-derived cardiomyocytes (hiPSC-CMs) were differentiated using a chemically defined monolayer differentiation protocol as previously described^11^. Briefly, iPSCs at 90% confluence were incubated with differentiation basal medium comprising RPMI 1640 medium (Thermo Fisher, 11875095) and B27 supplement minus insulin (Thermo Fisher, A1895601). CHIR99021 (8 μM) was added to the differentiation basal medium. On day 2, medium was removed and replaced with differentiation basal medium minus CHIR99021. On day 3, the Wnt antagonist, IWR-1, was added to the medium. After 48 hours, medium was removed and replaced with differentiation basal medium without any inhibitors. On day 7, the cells were incubated with complete CM medium consisting of RPMI 1640 medium and B27 supplement plus insulin (Thermo Fisher, 17504044). The medium was changed every 2 days. Monolayers of hiPSC-CMs were cultured for 30 days and subsequently dissociated for experimental use using TrypLE Express (Life Technologies). For measurement of inflammatory genes, hiPSC-CMs were exposed to either TNF-α (50 ng/mL) or EVs (control vs Nsp1) for 6 hours prior to RNA extraction. All procedures conformed to the UIC institutional review board-approved protocol.

### Production of lentivirus expressing SARS-CoV-2 subunits

Plasmids encoding for codon-optimized Nsp1, Nsp12, E and N of SARS-CoV-2 with a 2xStrep tag at the C-terminus were kindly provided by Dr. Nevan Krogan^8^. Empty vector or respective overexpression plasmids were packaged into virus using HEK293T cells as the packaging cell line in 10 cm dishes. Target DNA, helper plasmids VSVG and PAX2 were transfected at 9, 3 and 9 μg respectively using Lipofectamine 2000 (Thermo Fisher, 11668027). Infectious supernatant was collected at 48 and 72 hours after transfection and filtered to remove cell debris. Supernatant was then concentrated using Lenti-X Concentrator (TakaraBio, 631232) according to the manufacturer’s protocol. A549 cells were then transduced with either control or overexpression lentivirus with polybrene (8 μg/ml) overnight, and fresh medium supplemented with exosome-depleted FBS (Thermo Fisher, A2720803) was added and incubated for 48 hours before being collected for EV isolation.

### RNA extraction and quantitative real-time reverse transcription-polymerase chain reaction (qRT-PCR)

Total RNA was isolated using Direct-Zol RNA Miniprep Kit (Zymo Research, R2051). Reverse transcription was performed using the High-Capacity cDNA Reverse Transcription Kit with RNase Inhibitor (Thermo Fisher, 4374966) and qRT-PCR was performed using the PowerUp SYBR Green Master Mix (Thermo Fisher, A25742) on a QuantStudio 7 Flex real-time PCR detector (Thermo Fisher). Relative mRNA levels were normalized to those of GAPDH mRNA in each reaction and undetermined raw Ct values were set to 40 for analysis purposes. Primers sequences are shown in **Table 1**. Three to four replicates per group were used for qRT-PCR.

**Table 1.**
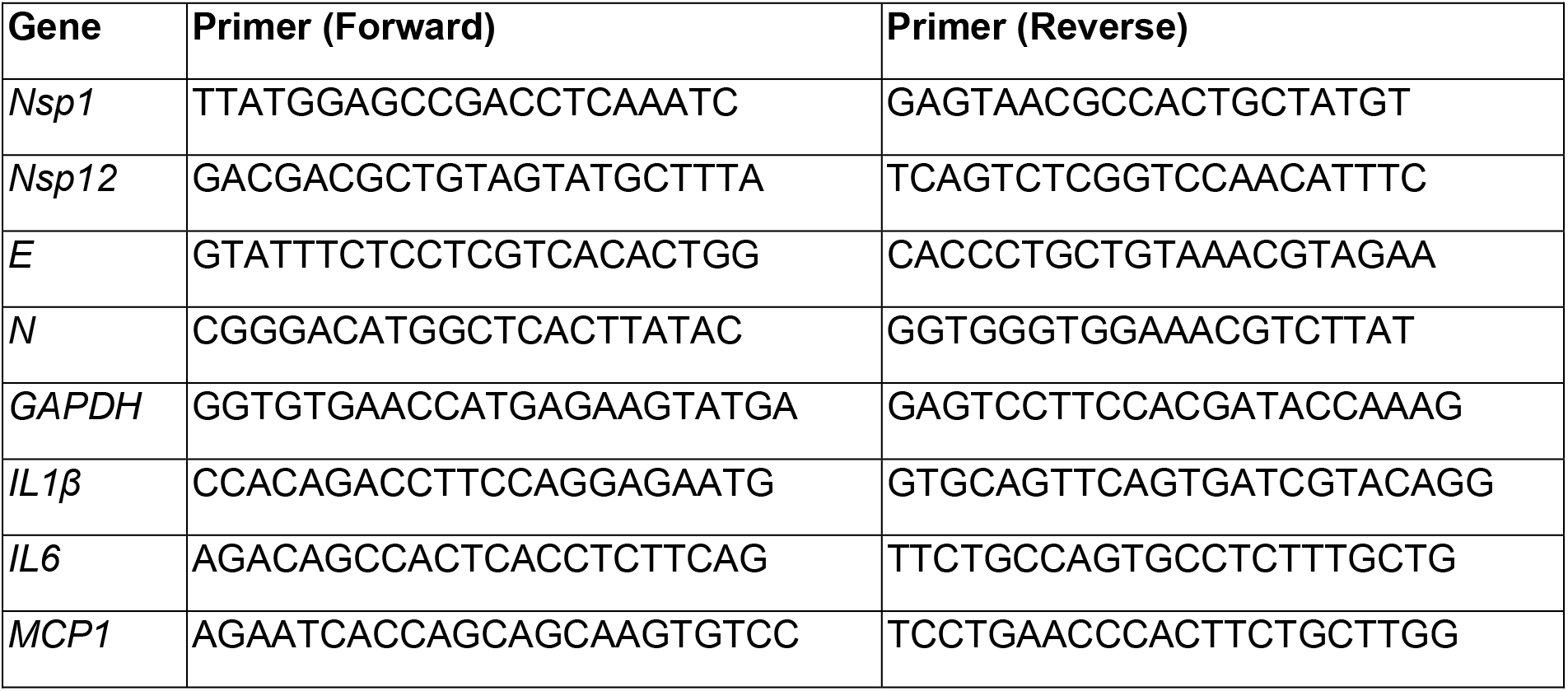
List of primers used in this study.

### Protein extraction and western blot analysis

EV samples were prepared in 1x RIPA buffer (Sigma, R0278) supplemented with protease and phosphatase inhibitor (Thermo Fisher, 78440). Samples were subjected to electrophoresis on 4-12% NuPAGE Bis–Tris gels (Thermo Fisher, NP0335BOX) and proteins were transferred to nitrocellulose membranes using wet-based transfer system (Bio-Rad). Membranes were incubated overnight with the indicated primary antibodies, followed by incubation for 1 hour with horseradish peroxidase-conjugated secondary antibodies (Cell Signaling; 7074 and 7076). Signals were detected by chemiluminescence. Primary antibodies used include CD63 (Thermo Fisher, 10628D), CD81 (Thermo Fisher, 10630D), TSG101 (Thermo Fisher, MA123296), calnexin (Thermo Fisher, MA3027) and albumin (Proteintech, 16475-1-AP).

### Isolation of extracellular vesicles

Supernatant of A549 cells was collected at 48 hours after transduction with lentivirus for isolation of EVs. Briefly, supernatant was first centrifuged at 300 x g for 5 minutes, followed by 1500 x g for 10 minutes, filtered (0.2 μM) and concentrated using Ultracel-100K (Millipore). EVs in concentrate were then isolated using Total Exosome Isolation Reagent (Thermo Fisher, 4478359) according to the manufacturer’s instructions overnight at 4°C, followed by centrifugation at 12 000 x g at 4°C for 1 hour. Isolated EVs were immunoblotted for EV markers CD63 and CD81. For immuno-magnetic isolation of CD63-positive EVs, the exosome-human CD63 isolation/detection reagent (Thermo Fisher, 10606D) was used based on the manufacturer’s protocol. EVs were pre-enriched using the Total Exosome Isolation Reagent as described above. For GW4869 (Sigma, D1692) experiments, supernatant was concentrated 100 times using Ultracel-100K following treatment of A549 cells for 48 hours with and without GW4869 (5 μM) before being added to hiPSC-CMs (1 x 10^6^ cells seeded in 1 mL media).

### RNase/protease treatment of EVs

An exosomal preparation was divided into four equal fractions for the following treatments: i) untreated; ii) RNase only, iii) RNase with proteinase K and iv) RNase, proteinase K and Triton X-100. First, 1% Triton X-100 or an equivalent volume of PBS was added and incubated on ice for 15 minutes. This was followed by the addition of proteinase K (0.05 μg/μl) at 37°C for 20 minutes, after which the reaction was stopped by adding 1x protease/phosphatase inhibitor cocktail. Finally, RNase A (0.5 μg/μl) is added at 37°C for 20 minutes. Samples were then immediately subjected to RNA isolation and qRT-PCR.

### Uptake of EVs by hiPSC-CMs

Purified EVs were labeled using the ExoGlow™-Protein EV Labeling Kit (System Bio, EXOGP400A-1) which labels internal EV proteins based on the manufacturer’s description. Briefly, 150 μg of EVs were resuspended in 500 μl of PBS and incubated with 1 μl of ExoGlow dye with shaking at 37°C for 20 minutes. For negative control staining, 500 ul of PBS + 1 μl of ExoGlow dye was used. Unincorporated dye was removed by the use of Exosome Spin Columns (Thermo Fisher, 4484449). Labeled EVs were then precipitated overnight as described above and added to hiPSC-CMs for 6 hours at 37°C. hiPSC-CMs were then fixed and stained with cardiac troponin T (Abcam, AB45932) and imaged by confocal microscopy to visualize uptake of labeled EVs.

### Statistics

The values presented are the means ± standard deviation from at least three samples. Statistical differences were determined by a 2-tailed, unpaired Student t-test or two-way ANOVA with Tukey’s multiple comparison test as appropriate. A value of P<0.05 was considered significant.

## Supporting information

Ct Values for Figures

## SUPPLEMENTAL FIGURE LEGENDS

**Supplemental Figure 1.**
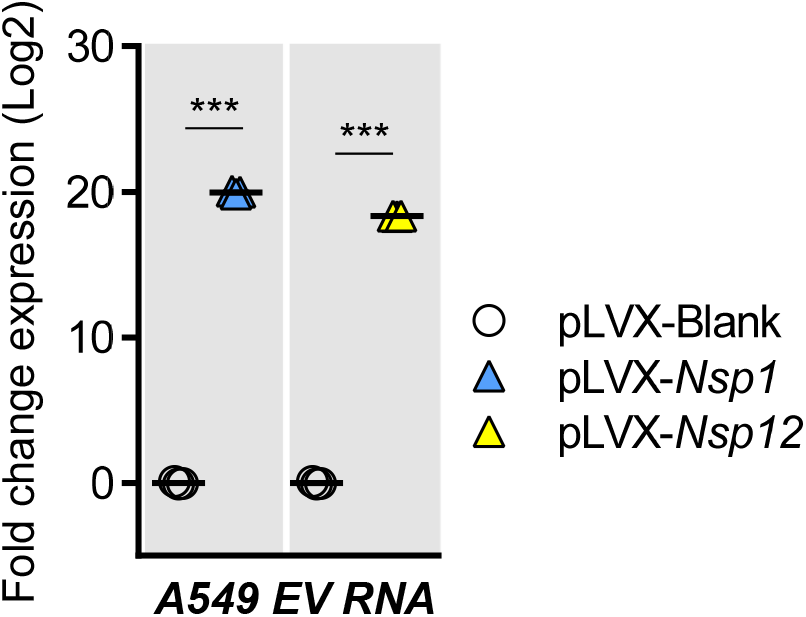
SARS-CoV-2 synthetic genes (Nsp1 and Nsp12) were detected in EVs released from A549 lung epithelial cells. EVs were isolated using immuno-magnetic anti-CD63 beads as an additional verification method. mRNA levels were measured by qRT-PCR (n = 3, mean ± S.D.). ***P<0.001 versus pLVX-Blank (Student’s t-test).

**Supplemental Figure 2.**
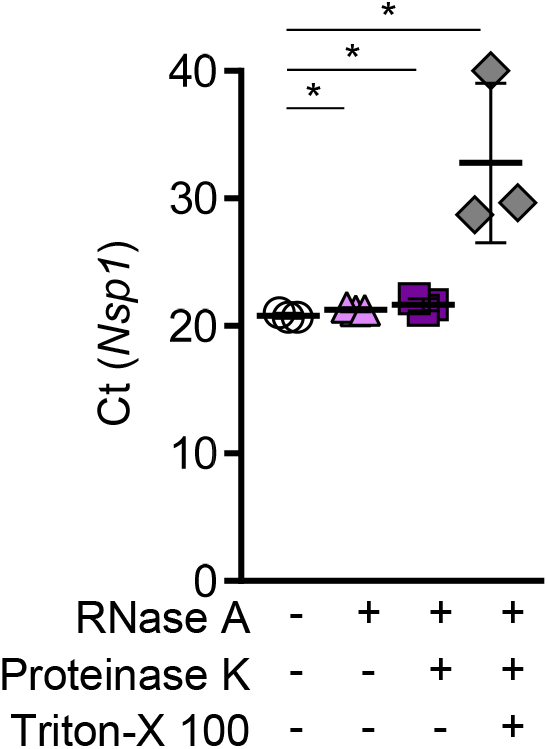
RNase A and Proteinase K sensitivity of Nsp1 within EVs. Nsp1 EVs were purified from A549 cell culture media and treated with either RNase A alone, RNase A and Proteinase K, or RNase A and Proteinase K in the presence of detergent. mRNA levels were measured by qRT-PCR (n = 3, mean ± S.D.). *P<0.05 versus untreated EVs (Student’s t-test).

**Supplemental Figure 3.**
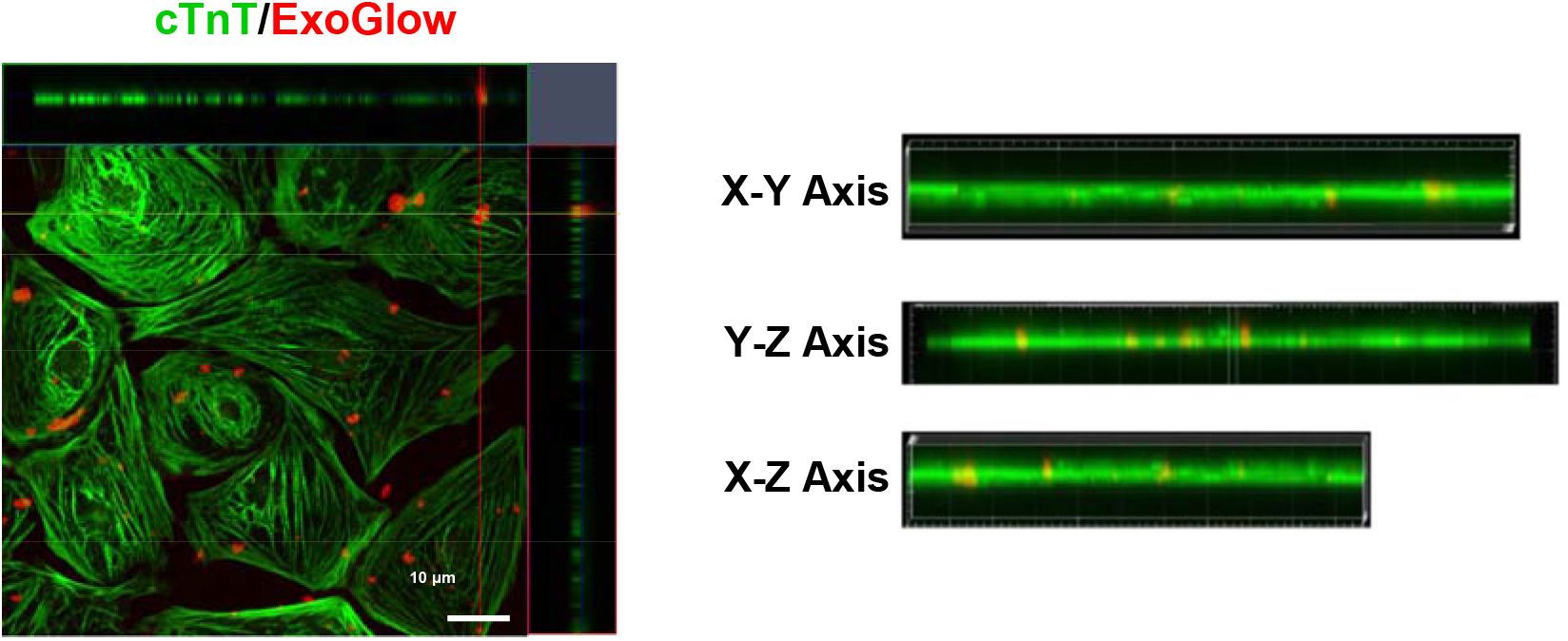
Visualization of ExoGlow-labeled exosomes (red) added to hiPSC-CMs which were stained with cardiac troponin T (green) by confocal imaging. Orthogonal planes (xy, yz and xz) of the confocal microscope images are depicted on the right. Scale bar = 10 μM.

**Supplemental Figure 4.**
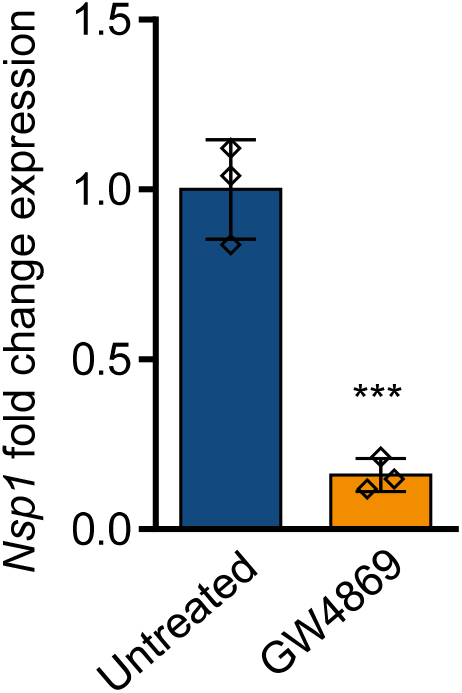
hiPSC-CMs were exposed to conditioned media collected from Nsp1-overexpressing A549 cells treated with or without GW4869 (5 μM). mRNA levels were measured by qRT-PCR (n = 3, mean ± S.D.). ***P<0.001 versus untreated (Student’s t-test).

